# Comparative Analysis of Neural Decoding Algorithms for Brain-Machine Interfaces

**DOI:** 10.1101/2024.12.05.627080

**Authors:** Olena Shevchenko, Sofiia Yeremeieva, Brokoslaw Laschowski

## Abstract

Accurate neural decoding of brain dynamics remains a significant and open challenge in brain-machine interfaces. While various signal processing, feature extraction, and classification algorithms have been proposed, a systematic comparison of these is lacking. Accordingly, here we conducted one of the largest comparative studies evaluating different combinations of state-of-the-art algorithms for motor neural decoding to find the optimal combination. We studied three signal processing methods (i.e., artifact subspace reconstruction, surface Laplacian filtering, and data normalization), four feature extractors (i.e., common spatial patterns, independent component analysis, short-time Fourier transform, and no feature extraction), and four machine learning classifiers (i.e., support vector machine, linear discriminant analysis, convolutional neural networks, and long short-term memory networks). Using a large-scale EEG dataset, we optimized each combination for individual subjects (i.e., resulting in 672 total experiments) and evaluated performance based on classification accuracy. We also compared the computational and memory storage requirements, which are important for real-time embedded computing. Our comparative analysis provides novel insights that help inform the design of next-generation neural decoding algorithms for brain-machine interfaces used to interact with and control robots and computers.

## I. Introduction

Brain-machine interfaces have enormous potential to allow patients with physical disabilities to interact with and control robots and computers by leveraging neuroimaging techniques such as electroencephalography (EEG). EEG can capture the neural dynamics from the brain associated with imagined limb movements, which are typically preceded by event-related desynchronization in the mu (8–12 Hz) and beta (18–26 Hz) rhythms, reflecting motor planning and execution. Following task completion or rest, these rhythms often exhibit event-related synchronization, making EEG a powerful tool for decoding neural signals [1]. However, accurately decoding brain dynamics into control commands remains a significant and open challenge. For example, EEG signals are prone to noise from eye movements and muscle activity [2]. Additionally, their low signal-to-noise ratio requires advanced signal processing and feature extraction to enhance data fidelity and extract meaningful neural information.

Different neural decoding algorithms for brain-machine interfaces have been proposed. For example, [3] developed a two-class neural decoder, comparing H∞ filters and artifact subspace reconstruction (ASR) with four different classifiers, including support vector machine (SVM), minimum distance to mean (MDM), MDM with filter, and linear discriminant analysis (LDA). In pseudo-online cross-validation experiments, they achieved accuracies of 66.2% for a generic model and 69.6% for a personalized model, with H∞ filter and MDM filter yielding the best neural decoding performance.

In terms of deep learning, [4] used recurrent long shortterm memory (LSTM) neural networks, comparing them to LDA classifiers. After a two-step signal preprocessing using ASR and independent component analysis (ICA), the authors extracted neural correlates from 1-40 Hz frequency bands. The LSTM neural network outperformed their LDA model during online evaluation, achieving accuracies of 83.3%, 80.5%, and 75.3%. Similarly, [5] compared common spatial patterns (CSP) with LDA and a convolutional neural network (CNN) classifier, finding the CNN significantly outperformed CSP with an LDA model by 15-18%.

Our review of the literature revealed a wide variety of different signal processing, feature extraction, and machine learning classification models used for motor neural decoding for brain-machine interfaces, with most studies creating their own individual algorithms. An important unanswered question is: “what is the optimal combination of these algorithms to decode neural dynamics from the brain”? To help answer this question, here we conducted one of the largest comparative studies to develop and test different neural decoding algorithms (i.e., consisting of 48 combinations of state-of-the-art signal processing, feature extraction, and machine learning classifiers, each optimized for individual subjects) using a large-scale EEG dataset to determine the optimal combination for brain-machine interfaces (Fig. 1).

**Fig. 1.**
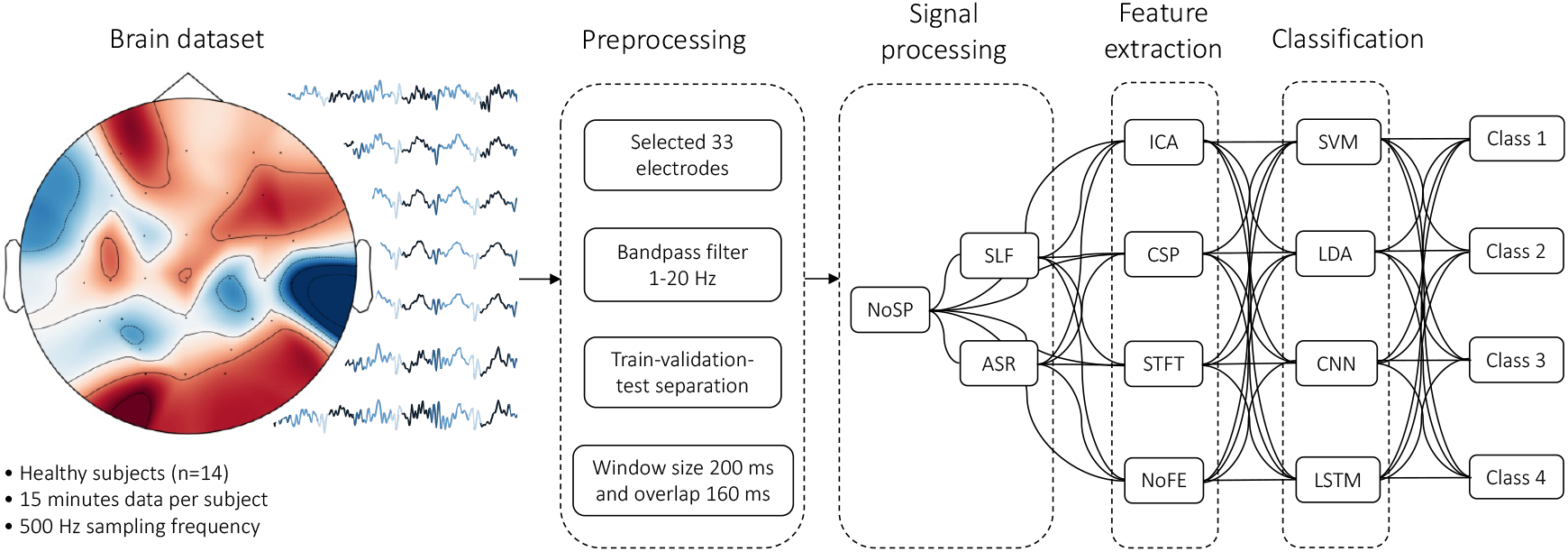
Our experimental design. We tested different state-of-the-art signal processing, feature extraction, and classification algorithms to find the optimal combination for decoding neural dynamics from the brain. Our baseline is no signal processing (NoSP) and no feature extraction (NoFE).

## II. Methods

### A. Brain Dataset

We used the open-source EEG dataset [6], which includes labels for gait events—”right heel strike” (190 ± 60 ms), “left heel strike” (180 ± 20 ms), “right toe off” (360 ± 20 ms), and “left toe off” (360 ± 20 ms)—from real-world walking experiments. We selected this dataset for its focus on short events, which require precise signal processing for accurate neural decoding. The brain data, recorded at 500 Hz from 26 healthy adults using 66 Ag/AgCl EEG electrodes in a custom 64-channel layout based on the 10-5 system, were referenced online to FCz. The foot movements were experimentally captured with 3D accelerometers, and EEG measurements were synchronized offline.

To maintain a large sample size while reducing experimental complexity, we used data from 14 subjects, with an average recording duration of 15 minutes per subject. We set aside the last two minutes of each recording as validation and testing sets to prevent data leakage. We kept the sampling rate at 500 Hz. We experimented with a number of different electrode subsets for EEG channel selection, including a 24-electrode set optimal for motor imagery and four custom sets targeting different cortical lobes in the brain. Although we found no statistically significant difference in F1-score, custom set, which included 33 electrodes, slightly outperformed the others with a 0.1 increase in F1-score.

For frequency filtering, we tested various combinations of high-pass (0.5–2.5 Hz) and low-pass (20–40 Hz) filters. We found that high-pass filters > 1.0 Hz significantly degraded classification, while low-pass filters > 20 Hz caused a slight reduction in accuracy. The optimal bandpass filter was 1–20 Hz, yielding the highest average F1-score across participants. Finally, we divided the data into windows, each labeled by the last element. We tested window lengths between 100-500 ms and overlaps of 20-160 ms. A window size of 200 ms with a 160 ms overlap provided one of the best F1-scores, generating fewer data points (i.e., 22,000 windows vs. 87,000 with a 150 ms window and 140 ms overlap) while still ensuring enough data for training without exceeding the smallest class size (Fig. 2). Fewer windows helped to reduce the memory storage and computing power requirements during signal processing.

**Fig. 2.**
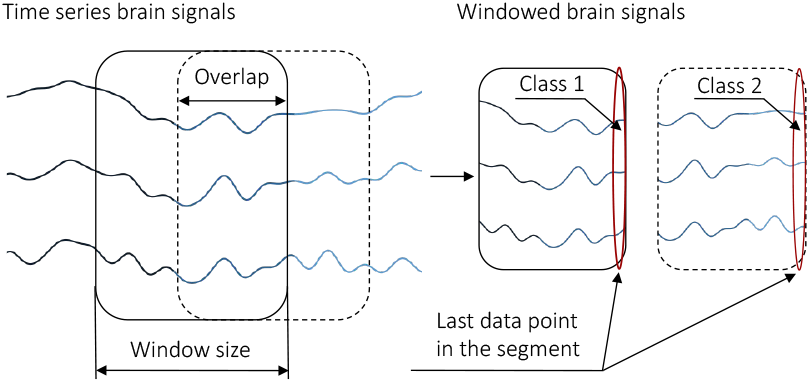
Schematic of our dataset adaptation used for semi-real-time neural decoding, including segmenting the brain signals with fixed window size and overlap. We used the class of the last data point in each segment to define the class of the entire segment.

Although each time series contains the same number of gait events, the duration of these events varies. Consequently, after sampling the data, as shown in Fig. 2, events with longer durations generated more windows, leading to an imbalanced final dataset. To address this, we evaluated two strategies: down sampling the overrepresented classes and retaining the original distribution. Up sampling was not considered due to the limited research on effective data augmentation techniques for EEG neural signals. Our analysis revealed that down sampling significantly reduced model performance. Therefore, we opted to retain the current class distributions to preserve the integrity of the neural data.

### B. Signal Processing

Common artifact removal methods in brain-machine interface decoding range from simple thresholding to more advanced methods like blind source separation [1], improving signal-to-noise ratio and data quality. In our study, we compared two different signal processing algorithms: 1) artifact subspace reconstruction (ASR), optimized to handle non-stationary artifacts, and 2) surface Laplacian filter (SLF), which increases local information by estimating the radial current flow at the scalp. We also used normalized data as our baseline, centering values around the mean with unit standard deviation.

*Artifact Subspace Reconstruction (ASR)* is an automatic method for removing transient, large-amplitude artifacts with minimal computational effort. It learns artifact characteristics directly from data, thus increasing specificity. While ASR cannot completely eliminate all artifacts—such as those from eye movements, muscle activity, or poor electrode contact—it can significantly improve data quality, facilitating further processing like ICA decomposition [7].

*Surface Laplacian Filter (SLF)*, or current source density, enhances neural data by emphasizing local brain dynamics and minimizing distant signal interference. It can improve spatial filtering, reduce noise, and highlight sensorimotor rhythms, by calculating the second spatial derivative of scalp potentials, enhancing topographical localization and reducing volume-conduction effects. Its effectiveness depends on electrode density and spacing, and it is independent of reference electrode choice. Positive estimates represent current flow from the brain to the scalp, while negative values indicate flow from the scalp to the brain [8].

### C. Feature Extraction

Feature extraction is important to transform complex brain dynamics into discriminative representations. By reducing data dimensionality, it highlights subtle neural modulations related to human movements, focusing on features such as eventrelated spectral perturbations and sensorimotor rhythms. This not only captures the spatial and temporal dynamics important for movement classification but also enhances the robustness and reliability of motor neural decoding.

In our study, we compared three state-of-the-art feature extraction methods used in brain-machine interfaces: common spatial patterns (CSP), which optimizes the spatial filters to distinguish brain dynamics; independent component analysis (ICA), which isolates the independent signals within the neural data; and short-time Fourier transform (STFT), which analyzes the frequency content of neural signals to capture both time and frequency information. We also used no feature extraction as our baseline for comparison.

*Common Spatial Patterns (CSP)*, initially developed for binary classification, decompose signals into spatial patterns that maximize variance between two states. For multi-class classification, we used the “one-versus-rest” method [9], generating spatial filters for each class against the rest. We factorized covariance matrices from each segment to extract eigenvectors corresponding to the largest eigenvalues, forming spatial filters to improve the signal-to-noise ratio and capture discriminative features critical for motor neural decoding [10].

*Independent Component Analysis (ICA)*, which is widely used in artifact removal and neural decoding, separates brain signals into independent sources. ICA finds an unmixing matrix based on component independence, enhancing the interpretability of brain data by isolating neural and non-neural sources. The independent components extracted serve as features, refining the representation of brain dynamics [11].

*Short-Time Fourier Transform (STFT)* applies Fourier transforms to windowed signal segments, providing time-localized frequency information. This method captures dynamic, time-specific brain activity, which is important for classifying event-related potentials, such as those associated with human movement. In our study, we used a semi-real-time decoding approach with windowed brain signals.

### D. Classification

The choice of classification algorithm is essential for motor neural decoding, influencing accuracy, computational efficiency, and real-time performance. We compared four different classifiers: two deep learning models (i.e., a convolutional neural network, CNN; and a long short-term memory network, LSTM) and two classical machine learning models (i.e., a support vector machine, SVM; and linear discriminant analysis, LDA).

*Support Vector Machine (SVM)* is a supervised machine learning algorithm that classifies data by finding the optimal hyperplane that maximizes the margin between classes, thus improving generalization. For non-linear classification, SVM uses a kernel function to map data into a higher-dimensional space where linear separation is possible.

*Linear Discriminant Analysis (LDA)* is a supervised method for classification and dimensionality reduction commonly used in neural decoding of brain dynamics [12]. The algorithm identifies a one-dimensional subspace such that a linear hyperplane best separates the classes while maximizing the difference in class means. It assigns larger weights to predictors that show greater class distinction.

*Convolutional Neural Network (CNN)* is a deep learning model that applies convolutions to capture spatial patterns. Increasingly used in brain-machine interfaces, CNNs are well-designed for neural decoding. In our experiments, we used EEGNet, which is the state-of-the-art CNN optimized for EEG neural decoding, which uses depthwise and separable convolutions to create an efficient deep learning model that captures key features while remaining lightweight [13].

*Long Short-Term Memory (LSTM)* networks are recurrent neural networks designed to process sequential data and capture temporal dynamics. We used the LSTM architecture by [4], which demonstrated over 80% accuracy in decoding human movements from brain signals. The network consists of two LSTM layers, where the output of the first layer serves as input to the second, followed by a fully-connected layer and a softmax activation function to return the class probabilities.

### E. Experiments

We designed a new pipeline to systematically compare the different signal processing, feature extraction, and classification algorithms for motor neural decoding. We conducted a total of 672 experiments, testing three signal processing methods, four feature extractors, and four machine learning classifiers, each of which were optimized for individual subjects. This resulted in 48 unique neural decoding algorithms per subject. Our experiment design is shown in Fig. 1. Our pipeline begins with preprocessing, including bandpass filtering (1–20 Hz), train-validation-test separation, and windowing with a fixed 200 ms rolling window size with a 160 ms overlap. The label “NoSP-NoFE” (i.e., no signal processing or feature extraction) served as our baseline for comparison.

Next, we applied the signal processing per subject. For ASR, we trained the model on clean calibration data with a K = 20 cutoff, applying it to the train, validation, and test segments. The result was labeled ASR-NoFE. For SLF, we used the spherical spline method, calculating G and H matrices based on electrode distances and Legendre polynomial terms. This produced three datasets for each subject: NoSP-NoFE, ASR-NoFE, and SLF-NoFE.

We then applied the different feature extractors. Our preliminary tests revealed that CSP features caused significant accuracy drops in multiclass classification, so we used projected EEG signals instead for classification. CSP failed to converge with SLF-processed data for four subjects, so we also excluded this combination. For ICA, we used the FastICA algorithm (max_iter = 2000 and tol = 0.0001) fitting the model with the calibration dataset and applying it to the segmented train, validation, and test datasets. For STFT, we applied Fourier transforms to each time window to obtain time-frequency representations for classifier input.

We then trained our four machine learning classifiers on each subject. We applied data normalization before the classification by standardizing each segment using the mean and standard deviation from the calibration train set. For SVM, we used the radial basis function kernel and optimized the regularization parameter C with cross-validation, testing values from 1 to 1000. For LDA, we used the shrinkage LDA, which regularizes class covariance matrices to prevent instability and overfitting. We selected the best shrinkage value via cross-validation, ranging from 0.01 to 1, using the least squares solver.

We trained our CNN model with hyperparameter optimization using four randomly selected subjects. We tuned batch size, number of temporal filters (F1), depth multiplier (D), and fully-connected (FC) size, with little variation in F1-scores (0.02). The final parameters were F1 = 8, D = 5, and batch size = 64. We also optimized the LSTM hyperparameters, although the results were less consistent than our CNN model. To address this, we used early stopping, scheduled learning rates, and fixed initial weights. The final parameters were batch size = 256, hidden size = 80, and fc size = 75. After training, we performed inference on the test set, calculating accuracy, precision, recall, and F1-score for each class, along with their weighted averages. Additionally, we measured compute time and memory usage. We carried out statistical comparisons using the Wilcoxon signed-rank test, with the null hypothesis that there is no significant difference in F1-scores between models. The NoSP-NoFE model served as our baseline. We ran all of our experiments on Google Colab with an NVIDIA T4 GPU and 50 GB of RAM.

## III. Results

Our results are summarized in Table 1. We used weighted F1-score because it is the most representative metric for multiclass classification and can capture misclassification. For all our machine learning classifiers, adding STFT feature extraction significantly reduced neural decoding performance by up to 25%. Among all subjects, the highest-performing neural decoders used the CNN classifier. Data processing appeared to have a modest effect across the different classifiers, but additional processing slightly reduced the performance variability compared to using only data normalization.

**Table 1.** Average F1-scores across all 14 subjects for each of our neural decoding algorithms, including different combinations signal processing (i.e., SLF and ASR), feature extraction (i.e., ICA and CSP), and machine learning classification (i.e., SVM, LDA, LSTM, and CNN). NoSP means no signal processing and NoFE means no feature extraction. The best combination(s) per classifier are bolded.

| Classifier | NoSP-NoFE | SLF-NoFE | ASR-NoFE | NoSP-ICA | NoSP-CSP | SLF-ICA | ASR-ICA | ASR-CSP |
| --- | --- | --- | --- | --- | --- | --- | --- | --- |
| SVM | 88.8±4 | 89.9±4 | 83.9±6 | 89.4±4 | 89.4±4 | 89.2±4 | 83.1±7 | 84.3±6 |
| LDA | 87.8±5 | 87.8±5 | 87.8±5 | 87.9±5 | 87.9±5 | 87.6±5 | 87.9±5 | 87.9±5 |
| LSTM | 78.7±10 | 82.4±6 | 82.2±9 | 78.9±10 | 86.3±10 | 78.8±10 | 83.9±8 | 81.0±15 |
| CNN | 91.9±3 | 91.7±3 | 91.2±3 | 91.8±3 | 91.8±3 | 92.9±3 | 91.0±3 | 90.9±3 |

For our SVM classifier, SLF produced the highest F1score, with a Wilcoxon test p-value of 0.0063, indicating a statistically significant improvement over our baseline model. ASR, in contrast, consistently decreased our neural decoding accuracy. The distribution of SVM results for each signal processing and feature extraction combination is shown in Fig. 3. For LDA, ICA typically produced the highest F1-score, but the Wilcoxon test (p-value = 0.3076) showed no statistically significant difference from our baseline. Fig. 4 shows the LDA results for each combination of signal processing and feature extraction.

**Fig. 3.**
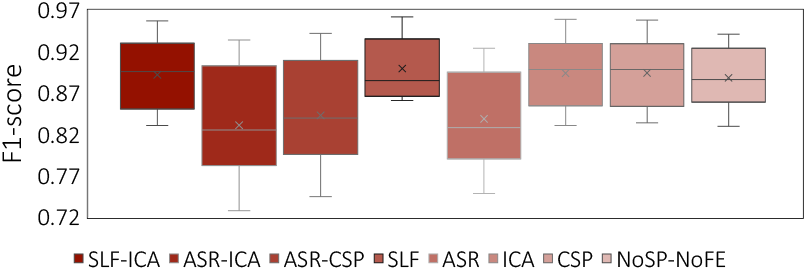
Neural decoding performance (i.e., weighted F1-score) using our support vector machine (SVM) classifier across all 14 subjects for each combination of signal processing and feature extraction.

**Fig. 4.**
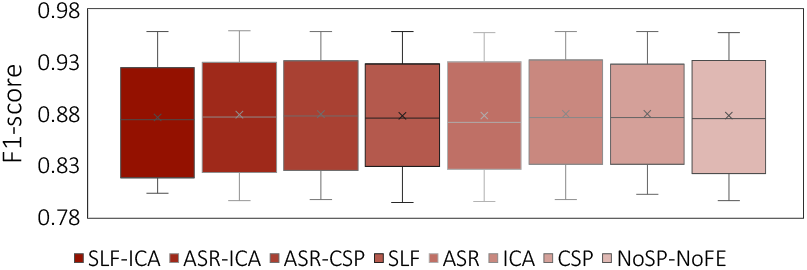
Neural decoding performance (i.e., weighted F1-score) using our linear discriminate analysis (LDA) classifier across all 14 subjects for each combination of signal processing and feature extraction.

For our LSTM classifier, CSP generally yielded the highest F1-score. The Wilcoxon test (p-value = 0.0001) confirmed a statistically significant improvement compared to our baseline. Fig. 5 shows the distribution of LSTM results across subjects. For our CNN classifier, the combination of SLF and ICA achieved the highest F1-scores, but the Wilcoxon test (p-value = 0.8597) showed no significant difference from our baseline. The CNN classification results are shown in Fig. 6. Tables 2-5 show several examples of confusion matrices using our CNN classifier for one subject.

**Table 2.**
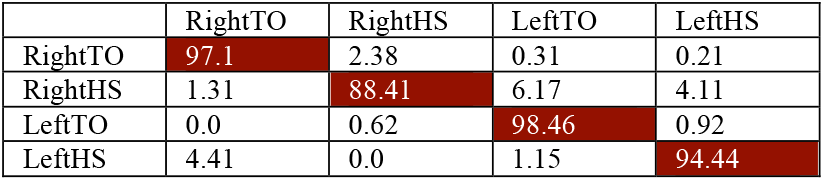
Example confusion matrix showing neural decoding accuracy (%) per class using our CNN model with NoSP and NoFE for one subject. The overall decoding accuracy was 95.5%.

**Table 3.** Example confusion matrix showing neural decoding accuracy (%) per class using our CNN model with SLF and NoFE for one subject. The overall decoding accuracy was 95.7%.

|  | RightTO | RightHS | LeftTO | LeftHS |
| --- | --- | --- | --- | --- |
| RightTO | 97.62 | 1.86 | 0.1 | 0.41 |
| RightHS | 2.24 | 88.79 | 4.67 | 4.3 |
| LeftTO | 0.0 | 1.13 | 98.15 | 0.72 |
| LeftHS | 3.26 | 0.38 | 1.72 | 94.64 |

**Table 4.** Example confusion matrix showing neural decoding accuracy (%) per class using our CNN model with NoSP and ICA for one subject. The overall decoding accuracy was 95.4%.

|  | RightTO | RightHS | LeftTO | LeftHS |
| --- | --- | --- | --- | --- |
| RightTO | 97.72 | 1.66 | 0.1 | 0.52 |
| RightHS | 2.06 | 87.66 | 5.98 | 4.3 |
| LeftTO | 0.1 | 1.13 | 98.25 | 0.51 |
| LeftHS | 3.64 | 0.38 | 2.3 | 93.68 |

**Table 5.** Example confusion matrix showing neural decoding accuracy (%) per class using our CNN model with SLF and ICA for one subject. The overall decoding accuracy was 95.4%.

|  | RightTO | RightHS | LeftTO | LeftHS |
| --- | --- | --- | --- | --- |
| RightTO | 97.72 | 1.76 | 0.1 | 0.41 |
| RightHS | 1.87 | 87.85 | 6.36 | 3.93 |
| LeftTO | 0.0 | 0.92 | 98.56 | 0.51 |
| LeftHS | 4.02 | 0.96 | 2.11 | 92.91 |

**Fig. 5.**
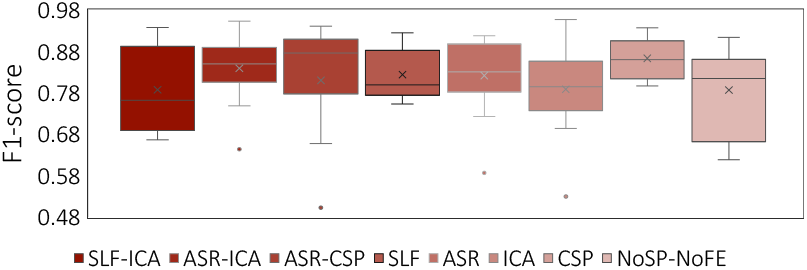
Neural decoding performance (i.e., weighted F1-score) using our long short-term memory (LSTM) neural network across all 14 subjects for each combination of signal processing and feature extraction.

**Fig. 6.**
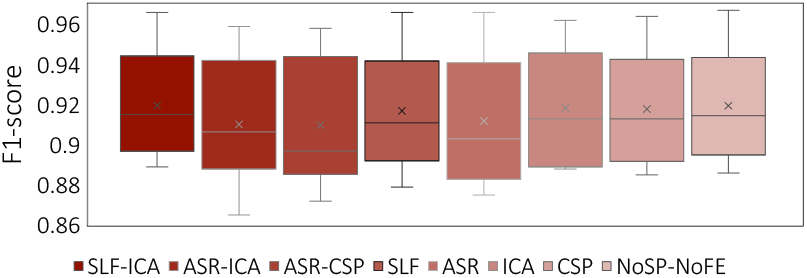
Neural decoding performance (i.e., weighted F1-score) using our convolutional neural network (CNN) classifier across all 14 subjects for each combination of signal processing and feature extraction.

In terms of other performance metrics, our model training took 2 minutes for CNN and LSTM, 2.5 minutes for LDA, and 6 minutes for SVM. Our smallest classification algorithm was the CNN model at 0.03 MB, while the LSTM model was 2 MB, and LDA and SVM models were 50.2 MB and 278.5 MB, respectively. Including signal processing and feature extraction increased processing times. The average time per data segment (100 data points, 33 features) was 4 ± 0.3 ms for ASR, 38 ± 3 ms for SLF, 0.3 ± 0.05 ms for ICA, 0.4 ± 0.06 ms for CSP, and 0.2 ± 0.05 ms for STFT.

## IV. Discussion

In this study, we conducted one of the largest comparative analyses testing different state-of-the-art signal processing, feature extraction, and classification algorithms to determine the optimal combination for motor neural decoding of brain dynamics. Using a large-scale EEG dataset of human movements in real-world environments, we tested three signal processing algorithms, four feature extractors, and four machine learning classifiers, each of which were optimized for individual subjects, resulting in 672 total experiments (i.e., 48 unique neural decoders per subject). We evaluated the performance of each combination based on weighted F1-score, inference speed, and model size, which have practical implications for real-time embedded computing. The results of our analyses can inform the design of next-generation neural decoding algorithms for brain-machine interfaces.

Our research showed that CNNs achieve the best neural decoding performance compared to classical machine learning algorithms and other deep learning models such as LSTMs. Our CNN significantly outperformed SVM, LDA, and LSTM models, with the highest F1-score observed when in combination with surface Laplacian filter (SLF) and independent component analysis (ICA) (93%). Although this improvement was not statistically significant compared to our baseline CNN model, it underscores the robustness of CNNs to different preprocessing. These results are consistent with previous research [5], which reported that CNNs optimized on raw EEG signals outperform the conventional CSP+LDA pipeline. Moreover, for neural decoding of human locomotion, we extended the application of LSTM models to four-class classification, exceeding the binary classification of previous research [4]. While [4] found that LSTM with ASR processing outperformed LDA with CSP, we observed the opposite. Our results showed that ASR processing increased performance variability and reduced decoding accuracy, suggesting that its utility requires careful evaluation. Our LSTM network achieved its best performance when combined with CSP signal processing, yielding statistically significant improvements over other combinations.

We also found that SVM achieves its best performance when paired with SLF processing, a statistically significant improvement over filtered and normalized data. These findings align with previous research [3], which found that SVM outperformed LDA under similar conditions. Furthermore, while our LDA classifier demonstrated its highest accuracy when combined with ICA, our Wilcoxon test revealed no significant difference compared to filtered and normalized data. These results highlight how classical models, despite their simplicity, can achieve competitive performance with appropriate signal processing. By comparing these models and signal processing algorithms, our study provides a comprehensive evaluation that bridges gaps in prior literature, offering deeper insights into the optimal neural decoder design.

Despite these developments, our study has several limitations. Our evaluation was limited to pseudo-online experiments aimed at developing and proving the feasibility of our decoding algorithms. However, brain-machine interfaces are real-time, closed-loop systems and thus eventually need to be tested online to demonstrate real-world performance. While we observed trends in higher F1-scores for some algorithms over others, we cannot be certain that these trends will generalize to other datasets or online measurements of brain dynamics. Furthermore, although the two signal processing and three feature extraction algorithms that we studied are among the most popular in motor neural decoding, especially for leg movements, other methods exist and should be considered in future research [3].

To address these limitations, we plan to extend the number of datasets and subjects used to optimize our neural decoding algorithms, including data of brain dynamics from more invasive recording technologies. Data from other sensors such as inertial sensors or egocentric vision from smart glasses could be integrated in a multimodal system together with brain data to improve accuracy in predicting human movements [14][20]. Furthermore, rather than optimizing neural decoders for individual subjects, as done in our study, future research could explore principles of transfer learning to develop a general model with the best combination of signal processing, feature extraction, and classification, trained on brain data from all subjects. This general model could then be compared to our neural decoding algorithms optimized for individual subjects.

In addition to other signal processing and feature extraction algorithms, we also plan to expand the number of machine learning classifiers for comparison such as hybrid CNNLSTM model, graph neural networks, unsupervised autoencoders and transformer-based models, which have recently gained significant attention. Several studies have already demonstrated that transformer-based models can outperform CNNs in EEG classification and prediction tasks [21]-[22]. A detailed comparison of these models using our new pipeline would be valuable. Combining multimodal signals in neural-machine interfaces could also offer more reliable input for diverse populations and applications [23]. We also plan to deploy our neural decoding algorithms to evaluate real-time embedded computing performance and extend our research to simulated environments [24]. Overall, we developed and tested dozens of neural decoding algorithms (i.e., consisting of different signal processing, feature extraction, and machine learning classifiers) using a large-scale EEG dataset to determine the optimal combination for brain-machine interfaces.

## Acknowledgment

This research study is dedicated to the people of Ukraine in response to Russia’s ongoing invasion.

## References

[1] R. Abiri et al., “A comprehensive review of EEG-based brain–computer interface paradigms,” Journal of Neural Engineering, Nov. 2018.

[2] I. Iturrate, R. Chavarriaga, and J. D. R. Millán, “General principles of machine learning for brain-computer interfacing,” Handbook of Clinical Neurology, vol. 168, pp. 311–328, Mar. 2020.

[3] V. Quiles et al., “Decoding of turning intention during walking based on EEG biomarkers,” Biosensors, vol. 12, pp. 555, Jul. 2022.

[4] S. Tortora et al., “Deep learning-based BCI for gait decoding from EEG with LSTM recurrent neural network,” Journal of Neural Engineering, vol. 17, pp. 046011, Jul. 2020.

[5] N. Tibrewal, N. Leeuwis, and M. Alimardani, “Classification of motor imagery EEG using deep learning increases performance in inefficient BCI users,” Plos One, vol. 17, pp. e0268880, Jul. 2022.

[6] N. S. J. Jacobsen, S. Blum, K. Witt, and S. Debener, “A walk in the park? Characterizing gait-related artifacts in mobile EEG recordings,” European Journal of Neuroscience, vol. 54, pp. 8421–8440, Dec. 2021.

[7] C. Y. Chang, S. H. Hsu, L. Pion-Tonachini, and T. P. Jung, “Evaluation of artifact subspace reconstruction for automatic artifact components removal in multi-channel EEG recordings,” IEEE Transactions on Biomedical Engineering, vol. 67, pp. 1114–1121, Jul. 2019.

[8] S. Deng, W. Winter, S. Thorpe, and R. Srinivasan, “Improved surface Laplacian estimates of cortical potential using realistic models of head geometry,” IEEE Transactions on Biomedical Engineering, vol. 59, pp. 2979–2985, Nov. 2012.

[9] Z. Y. Chin, K. K. Ang, C. Wang, C. Guan, and H. Zhang, “Multi-class filter bank common spatial pattern for four-class motor imagery BCI,” Annual International Conference of the IEEE Engineering in Medicine and Biology Society, Nov. 2009.

[10] H. Ramoser, J. Muller-Gerking, and G. Pfurtscheller, “Optimal spatial filtering of single trial EEG during imagined hand movement,” IEEE Transactions on Rehabilitation Engineering, vol. 8, pp. 441–446, Dec. 2000.

[11] A. S. Oliveira, B. R. Schlink, W. D. Hairston, P. König, and D. P. Ferris, “A channel rejection method for attenuating motion-related artifacts in EEG recordings during walking,” Frontiers in Neuroscience, vol. 11, pp. 225, Apr. 2017.

[12] D. Liu et al., “EEG-based lower-limb movement onset decoding: Continuous classification and asynchronous detection,” IEEE Transactions on Neural Systems and Rehabilitation Engineering, vol. 26, pp. 1626–1635, Aug. 2018.

[13] V. J. Lawhern et al., “EEGNet: a compact convolutional neural network for EEG-based brain–computer interfaces,” Journal of Neural Engineering, vol. 15, pp. 056013, Jul. 2018.

[14] O. Tsepa, R. Burakov, B. Laschowski, and A. Mihailidis, “Continuous Prediction of Leg Kinematics during Walking using Inertial Sensors, Smart Glasses, and Embedded Computing,” IEEE International Conference on Robotics and Automation (ICRA), May. 2023.

[15] H. Tan, A. Mihailidis, and B. Laschowski, “Egocentric perception of walking environments using an interactive vision-language system,” bioRxiv, 2024.

[16] D. Kuzmenko, O. Tsepa, A. G. Kurbis, A. Mihailidis, and B. Laschowski, “Efficient visual perception of human-robot walking environments using semi-supervised learning,” IEEE International Conference on Intelligent Robots and Systems (IROS). Dec. 2023.

[17] B. Ivanyuk-Skulskiy, A. G. Kurbis, A. Mihailidis, and B. Laschowski, “Sequential image classification of human-robot walking environments using temporal neural networks,” IEEE International Conference for Biomedical Robotics and Biomechatronics (BioRob), Oct. 2024.

[18] A. G. Kurbis, D. Kuzmenko, B. Ivanyuk-Skulskiy, A. Mihailidis, and B. Laschowski, “StairNet: Visual recognition of stairs for human-robot locomotion,” BioMedical Engineering OnLine, Feb. 2024.

[19] D. Rossos, A. Mihailidis, and B. Laschowski, “AI-powered smart glasses for sensing and recognition of human-robot walking environments,” IEEE International Conference for Biomedical Robotics and Biomechatronics (BioRob), Sept. 2024.

[20] D. Rossos, A. Mihailidis, and B. Laschowski, “AI-powered smart glasses for sensing and recognition of human-robot walking environments,” IEEE International Conference for Biomedical Robotics and Biomechatronics (BioRob), Sept. 2024.

[21] W. Cui et al., “Neuro-GPT: Towards a foundation model for EEG,” arXiv, Mar. 2024.

[22] Y. Song, X. Jia, L. Yang, and L. Xie, “Transformer-based spatialtemporal feature learning for EEG decoding,” arXiv, Jun. 2021.

[23] J. Thomas and B. Laschowski, “Development of a real-time neural controller using an EMG-driven musculoskeletal model,” bioRxiv, 2024.

[24] A. Dashkovets and B. Laschowski, “Reinforcement learning for control of human locomotion in simulation,” IEEE International Conference on Biomedical Robotics and Biomechatronics (BioRob), Oct. 2024.

